# A higher-order equivalence of Lotka-Volterra and replicator dynamics reveals tight connections between ecology and evolution

**DOI:** 10.1101/2025.03.28.645916

**Authors:** Chaitanya S. Gokhale, Arne Traulsen

**Author notes:** These authors contributed equally to this work.

## Abstract

The Lotka-Volterra equations are foundational in ecology, modelling logistic growth in isolated populations and complex dynamics in interacting species. Hofbauer and Sigmund established that these equations are equivalent to the replicator dynamics of evolutionary games, so dynamic patterns in ecology are mirrored in evolutionary games and vice versa. Both fields have since turned to non-linearities and higher-order interactions, where it is unclear whether this equivalence still holds. Here, we demonstrate the general equivalence and illustrate it in classical non-linear models from theoretical ecology. Non-linearities in either field leave this foundational connection intact. Yet directly modelling ecological dynamics with evolutionary games, or the reverse, risks misinterpretation, since species in Lotka-Volterra dynamics cannot be equated with strategies in evolutionary games. Our study reveals the tight interplay between ecology and evolutionary game theory, and the robustness of their mathematical connection even in complex scenarios, alongside its caveats.

---

The Lotka-Volterra equations represent the most fundamental model in ecology, describing the growth of interacting species within a population. While these equations result in logistic growth for isolated populations, interactions between species can lead to complex dynamics, including limit cycles and chaos. This wide applicability and the potential to capture some of the most fundamental characteristics of living beings, interactions, has sustained researchers’ interest in the model theoretically and empirically (*1–3*).

Another set of equations - the replicator equations - shares the spotlight due to their versatility of interpretations, ranging from Fisher’s selection equation to Bayesian updating (*4–8*). Hofbauer and Sigmund demonstrated an equivalence between the Lotka-Volterra equations and evolutionary game theory, showing that the dynamics of *n* species in Lotka-Volterra are dynamically equivalent to the replicator dynamics of an *n* + 1 strategy matrix game (*9, 10*). If we find a specific dynamic pattern in one of these systems, e.g., a stable limit cycle, we will find the same dynamic pattern in the corresponding system. For instance, the classic predator-prey dynamics are analogous to a Rock-Paper-Scissors game (*11*). However, with this transformation the equivalence between species and strategies becomes ambiguous, especially with the additional dimension. The extension of the dimension from *n* to *n* + 1 in the transformation goes back to a lost degree of freedom: In the Lotka-Volterra equations, population sizes can freely vary, whereas in replicator dynamics, relative abundances add up to one. Therefore, the elegant mathematical transformation of Hofbauer and Sigmund leads to a conceptual challenge, as the equivalence between species and strategies is muddled, and it is unclear how the additional type has to be interpreted in any specific biological system (*12*).

An important extension of the Lotka-Volterra equations in theoretical ecology involves considering higher-order interactions, in which the presence or absence of one species affects interactions among the others. This complexity necessitates more advanced models, often incorporating higher-order polynomials (*13–15*). Such non-linear interactions have recently gained increased attention, with much of the analysis relying on complex simulations and numerical methods (*14, 16*) as analytical advances are often challenging (*15, 17*).

A common critique of traditional interaction models is their lack of biological realism - e.g., growth rates do not immediately increase with the availability of more prey, but can be influenced by factors like handling times (*18*). Additionally, group size is a well-documented source of non-linearity in ecological interactions. Animal group sizes, from sparrows to humans, play a critical role in both intra- and interspecies interactions (*19,20*). For example, in the mutualistic relationship between Lycaenid larvae and ants, where ants protect the larvae in exchange for honeydew, the nutritional contribution of the larvae decreases when they are grouped with others rather than being solitary (*21, 22*). Moreover, this contribution depends on the number of ants attending each larva, highlighting the importance of group sizes in determining the costs and benefits of interspecies interactions. Predator-prey dynamics, a classical mainstay of ecological theory, are often much more complicated in practice. Group hunts in large mammals, from lions to wolves to killer whales, are often a function of the number of predators, their roles in the hunt, the herd size of the prey, and the group’s defence capabilities (*23–25*). Such complications in predator-prey studies, and similarly in host-parasite interactions, often naturally lead to non-linearities even in simple two-species dynamics (*26,27*). As a result, ecological models often require more sophisticated functional responses that account for these non-linearities.

In parallel, evolutionary game theorists have explored multiplayer games, where complex dynamics can arise even with a few interacting strategies (*28–34*). A classic example is the “tragedy of the commons”, which can be extended beyond linear, pairwise interactions to capture the non-linear dynamics of biological populations (*35–37*). As the replicator dynamics for multiplayer games develops further, a tight mathematical connection between these and ecological models has so far remained elusive.

Here, we demonstrate that the equivalence between ecological models and evolutionary games persists even for higher-order interactions. Although the equivalence follows Hofbauer and Sigmund’s framework, higher-order interactions complicate the notation and interpretation, and the interaction structures increase in dimensionality. We underscore the fundamental nature of this high-dimensional equivalence using some paradigmatic ecological non-linear models – the Holling functional response and the Allee effect. Moreover, the existing conceptual challenges - such as the distinction between species in ecological models and strategies in game theory - are amplified, necessitating careful consideration when applying these models to real-world biological systems (*12*).

## Established and establishing equivalence

### Pairwise interactions

The Lotka-Volterra and replicator equations are two mathematical models describing the dynamics of interacting populations or strategies in biology and game theory. They are related by a transformation that preserves some properties of the systems, such as the existence and stability of equilibria, but not the time scale of the dynamics. First, we recall the equivalence between the two models.

Lotka and Volterra independently derived equations for chemical and ecological systems (*38,39*) that, in their general form, are given by,

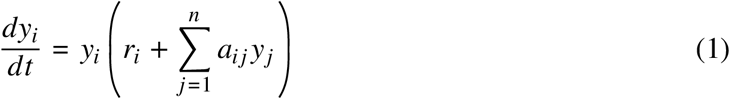

where *y*_*i*_ is the population size or density of species *i, r*_*i*_ is its intrinsic growth rate, and *a*_*ij*_ is the interaction coefficient between species *i* and *j*. There are *n* growth rates *r*_*i*_, one for each species.

The *n* × *n* matrix *A* = *a*_*ij*_ captures the effects of competition, predation, parasitism, or mutualism between and within the *n* different species (Fig. 1).

**Figure 1:**
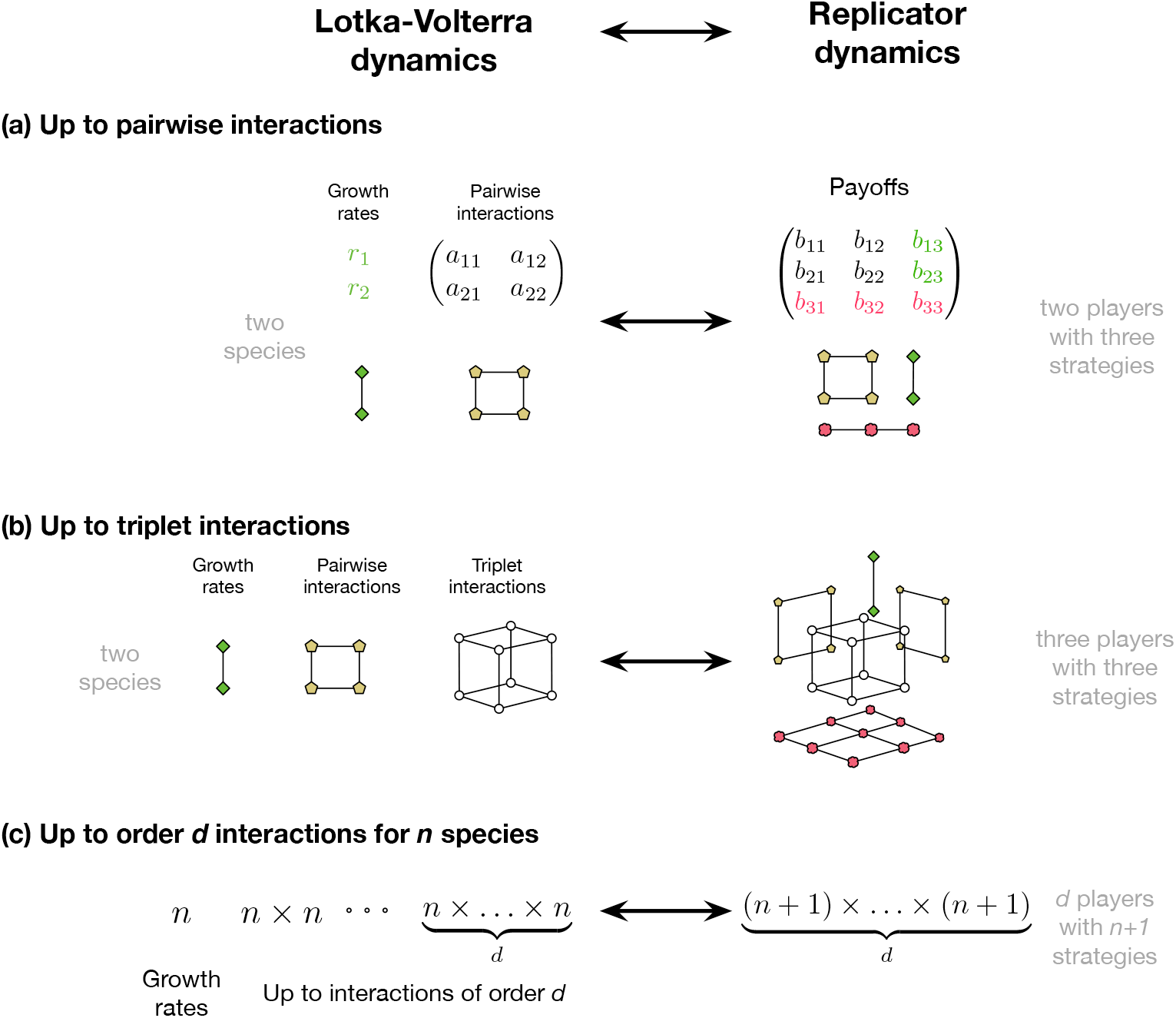
Establishing equivalence between Lotka-Volterra evolutionary games for higher-order interactions. The Lotka-Volterra dynamics include the growth rates and (inter-intra) species interactions as parameters. These parameters can be subsumed into the payoff matrix of an evolutionary game with equivalent dynamics. **(a)** Equivalence between pairwise interactions (yellow-green pentagons) between species and the replicator dynamics for two-player games is shown, where the growth rates from LV appear in the payoff matrix (green diamonds). The *n* species dynamics translates to a *n* + 1 strategy space where the payoff entries for the new strategy are all 0s (denoted in red, cloudy). **(b)** Including triplet interactions (cube with circular vertices) in LV dynamics translates to a three-player evolutionary game. Again, we can position the growth rates and the extra strategy payoffs in the tensor that already includes the growth rates (green diamonds), pairwise interactions (yellowish-green pentagons), and the third strategy payoffs, which are all 0s (red clouds). **(c)** To convert an LV system with all possible orders on interactions from order zero (growth rate) to *d*^th^ order, we make use of a *d*-dimensional tensor of size (*n* + 1)^*d*^ to capture all the payoff entries.

The term “replicator dynamics” was coined in 1983 (*4*) for a set of equations introduced slightly earlier (*40, 41*). The replicator equation has become seminal in the analysis of evolutionary games (*42*) where the dynamics of *n* + 1 strategies follow,

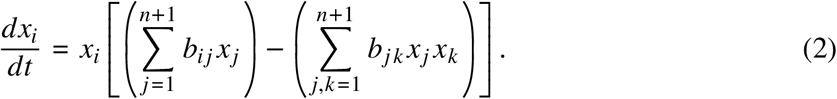

Here, *x*_*i*_ is the relative abundance of strategy *i* (such that 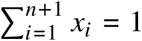), *b*_*ij*_ is the payoff to strategy *i* in a population when interacting with another strategy of type *j*. The second term captures the average payoff (*43*).

Hofbauer and Sigmund connected evolutionary game theory to theoretical ecology by mapping the replicator dynamics to the Lotka-Volterra equations (*10*). A set of Lotka-Volterra equations for *n* species can be transformed into a set of replicator equations for *n* + 1 strategies. For this transformation, the *n* × *n* species interaction matrix has to be cast into an (*n* + 1) × (*n* + 1) payoff matrix used to calculate the average fitnesses of the strategies in the replicator dynamics. The extended payoff matrix subsumes the *n* different intrinsic growth rates *r*_*i*_. The transformation,reiterated in Appendix), preserves the dynamical properties of the systems, such as the existence and stability of equilibria or limit cycles.

An example of the transformation is shown in Fig. 2. The Lotka-Volterra dynamics show the two species populations spiralling towards a fixed point. A transformed replicator model shows a similar spiral towards a coexistence equilibrium between three strategies (code for generating both figures is available online).

**Figure 2:**
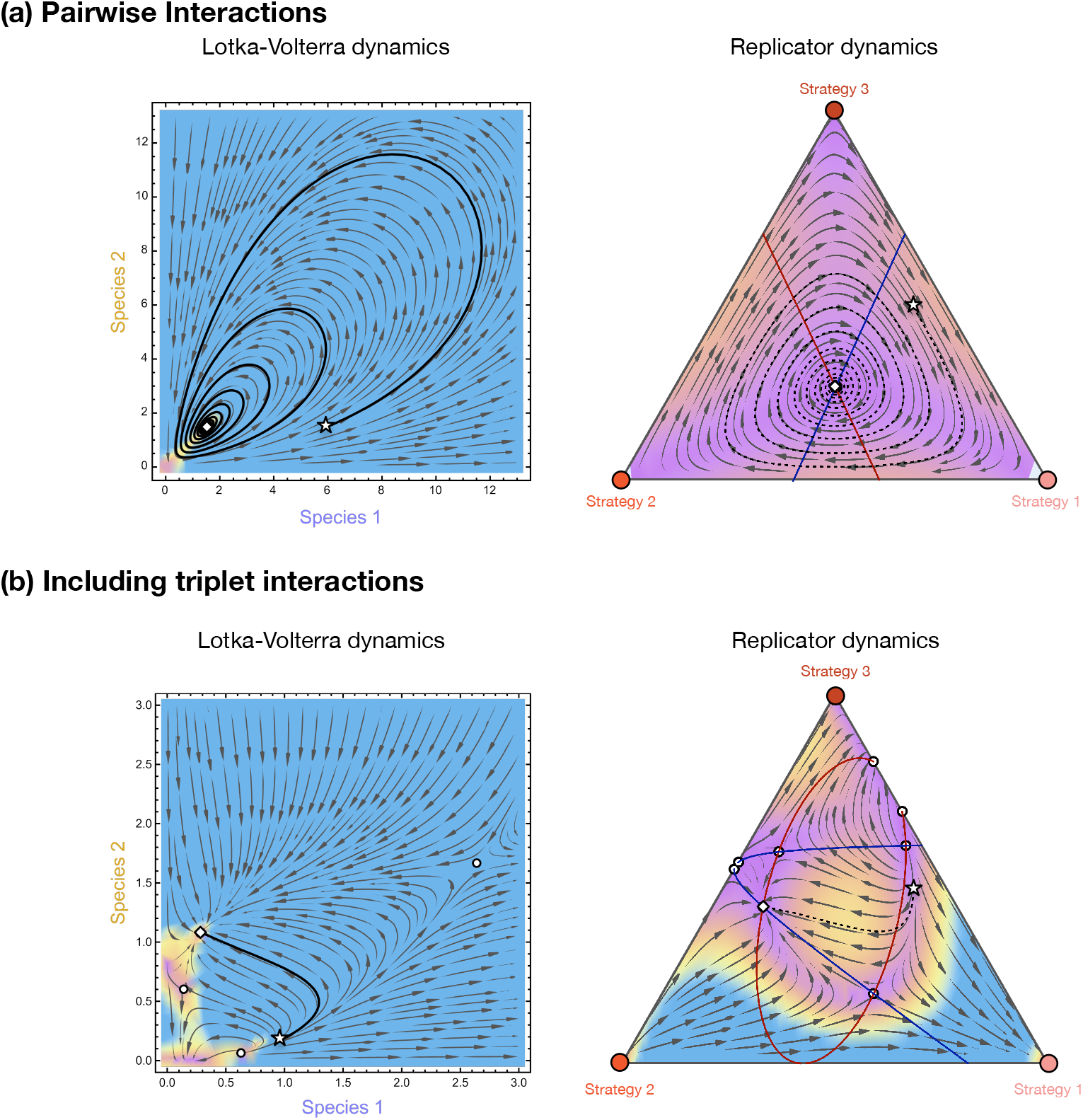
Equivalence between the Lotka-Volterra dynamics and the Replicator Dynamics. **(a)** As an example for pairwise interactions in the Lotka-Volterra system, quadratic in nature, we show the dynamics starting from an initial point (⭐) and spiralling towards a fixed point (⋄). The equivalent replicator dynamics for three types, cubic in nature, is represented here by the simplex on the right – spiraling towards the equivalent internal fixed point. **(b)** A three-player evolutionary game with three strategies can show a maximum of 2^2^ = 4 internal fixed points as it is a quartic equation. As an example, we show a representative numerical trajectory that converges to an internal stable fixed point. When converted to an equivalent Lotka-Volterra system, to a cubic equation, numerical integration yields an internal fixed point. Colored lines show nullclines of the dynamics. The code for generating the figures is available on Zenodo.

### Equivalence in higher dimensions

Having reiterated the established connection between (1) and (2), we now move to the following Lotka Volterra system that captures up to second order interactions as

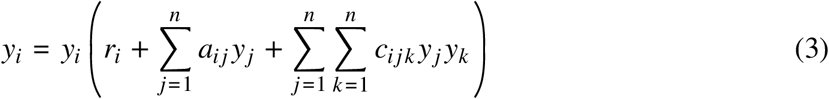

where *r*_*i*_ represent the growth rate of species *i* independent of any interactions, *a*_*ij*_ represents the impact of pairwise interactions of *j* on *i*, and *c*_*ijk*_ represents the joint impact of species *j* and *k* on *i*. Note that the indices *i, j* and *k* can be the same; for example, −*r*_*i*_/ *a*_*ii*_ can be interpreted as the carrying capacity of species *i* in isolation and *c*_*iii*_ corresponds to an allele effect of species *i* in isolation. In *c*_*ijk*_, the different index locations can be interpreted as the order in which the species interact (*14*). Eq. (3) naturally takes into account such complexities as it combines pairwise interactions (*a*_*ij*_) with triplet interactions (*c*_*ijk*_) as in (*15*). Higher-order interactions do not imply more than two species (*44, 45*): The additional term *c*_*ijk*_ does not affect the number of species *n*, but leads to non-linear interactions between them.

Converting the Lotka-Volterra system from Eqs. (3) to a replicator system with only pairwise interactions is impossible. One natural way to introduce higher-order interactions in evolutionary games is through multiplayer games. Going from an *n*-dimensional second-order Lotka-Volterra system to an *n* + 1-dimensional three-player replicator system necessitates constructing a payoff tensor of size (*n* + 1) × (*n* + 1) × (*n* + 1). For example, a three-player game with three strategies can be captured by a tensor **B** = *b*_*ijk*_ of size 3 × 3 × 3. (Fig. 1 middle right). In Appendix, we show how the mapping works. There is a clear algorithm on how to construct *b*_*ijk*_: This payoff tensor has a structure that emerges from the algorithm, transforming the interaction structures and the growth rates (see Appendix.). We start with setting *b*_*ijk*_ = *c*_*ijk*_ for *i, j, k* = 1, …, *n*. Next, we include the pairwise interactions into the tensor at *b*_*i,n*+1,*k*_ = *a*_*i,j*_. Further, we include the intrinsic growth rates *b*_*i,n*+1,*n*+1_ = *r*_*i*_. Finally, all other entries are set to 0 (Fig. 1). This construction reflects a many-to-one feature of the mapping. Any choice of (*b*_*i,n*+1,*k*_, *b*_*i,k,n*+1_)satisfying *b*_*i,n*+1,*k*_ + *b*_*i,k,n*+1_ = *a*_*i,k*_ yields the same Lotka-Volterra dynamics. We have arbitrarily set *b*_*i,k,n*+1_ = 0 as a canonical choice, but alternative splits correspond to genuinely different multiplayer games, in particular, games in which the order of action between players matters, capturing priority effects (*14*). These games are equivalent at the level of the deterministic ecological dynamics, but they can differ in their interpretation as games and in their behaviour under finite-population stochastic dynamics or in spatial settings. The deterministic LV-to-game map is therefore one-to-many, with the family of equivalent games related by redistributing weight among symmetric tensor entries.

With this, we can construct a system of replicator equations equivalent to Eqs. 3,

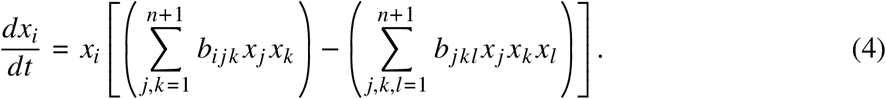

Given an evolutionary game, the payoff structure is not always in the form amenable to a Lotka-Volterra transformation. However, it is possible to move from any given evolutionary multiplayer game to an equivalent Lotka-Volterra system. In a three-player game, the payoff *f*_*i*_ of strategy *i* is given by *f*_*i*_ = ∑_*j,k*_ *b*_*ijk*_ *x*_*j*_ *x*_*k*_. The strategy dynamics are then governed by the difference between *f*_*i*_ and the average fitness 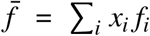. In a two-player game, the system’s dynamics are invariant to adding arbitrary constants to the columns of the payoff matrix (*11*). Similarly, for the tensor **B**, the property on the invariance of the dynamics to the addition of a constant to a plane still remains true. Addition of a constant to a plane e.g. *b*_*ijk*_ → *b*_*ijk*_ + Γ_*jk*_ results in *x*_*i*_ = *x*_*i*_ (∑_*j,k*_ (*b*_*ijk*_ + Γ_*jk*_*) x*_*j*_ *x*_*k*_ –(∑_*j,k,l*_ *x*_*j*_ (*b*_*jkl*_ + Γ_*kl*_ *)x*_*k*_ *x*_*l*_) equivalent to the replicator dynamics given by *x*_*i*_ = *x*_*i*_ (∑_*j,k*_ *b*_*ijk*_ *x*_*j*_ *x*_*k*_ – (∑_*j,k,l*_ *x*_*j*_ *b*_*jkl*_ *x*_*k*_ *x*_*l*_)). Similarly, multiplying a positive constant by a plane will only alter the speed of the dynamics but not the qualitative outcome. We follow the approach of (*10*) but adjust it to our purpose. An example of the transformation is shown in Fig. 2 (bottom). Here, we start with a three player game and convert it to an equivalent Lotka-Volterra system (code and transformation provided online).

In Appendix, we show that a complementary approach holds for any higher-order interaction of degree *d*. For example, two species dynamics with interactions of quartic order can be mapped into a three strategy game with four-player-interactions (Fig. 2 bottom).

### Specific models - from ecology to evolutionary games

So far, we have introduced the conversion between models from evolutionary games to ecology, and back, in abstract terms. Now, we use concrete ecological models that already include non-linearities to demonstrate the mathematical equivalence in action. Some of the most fundamental models of ecological interactions are already non-linear.

#### Holling’s functional responses

A classic ecological model is the case of predator handling exemplified by Holling (*18*). Holling’s type II functional response highlights the importance of the time required to process food. Hence, the consumption rate saturates at a certain level even as prey density continues to increase. To capture these non-linear response curves, we need non-linear interaction terms in the dynamics (See App.). Translating these ecological models to evolutionary games is possible with the equivalence we explained above. We show the equivalence with a three-player three-strategy evolutionary game by approximating the type II functional response for short interaction times in Figure 3.

**Figure 3:**
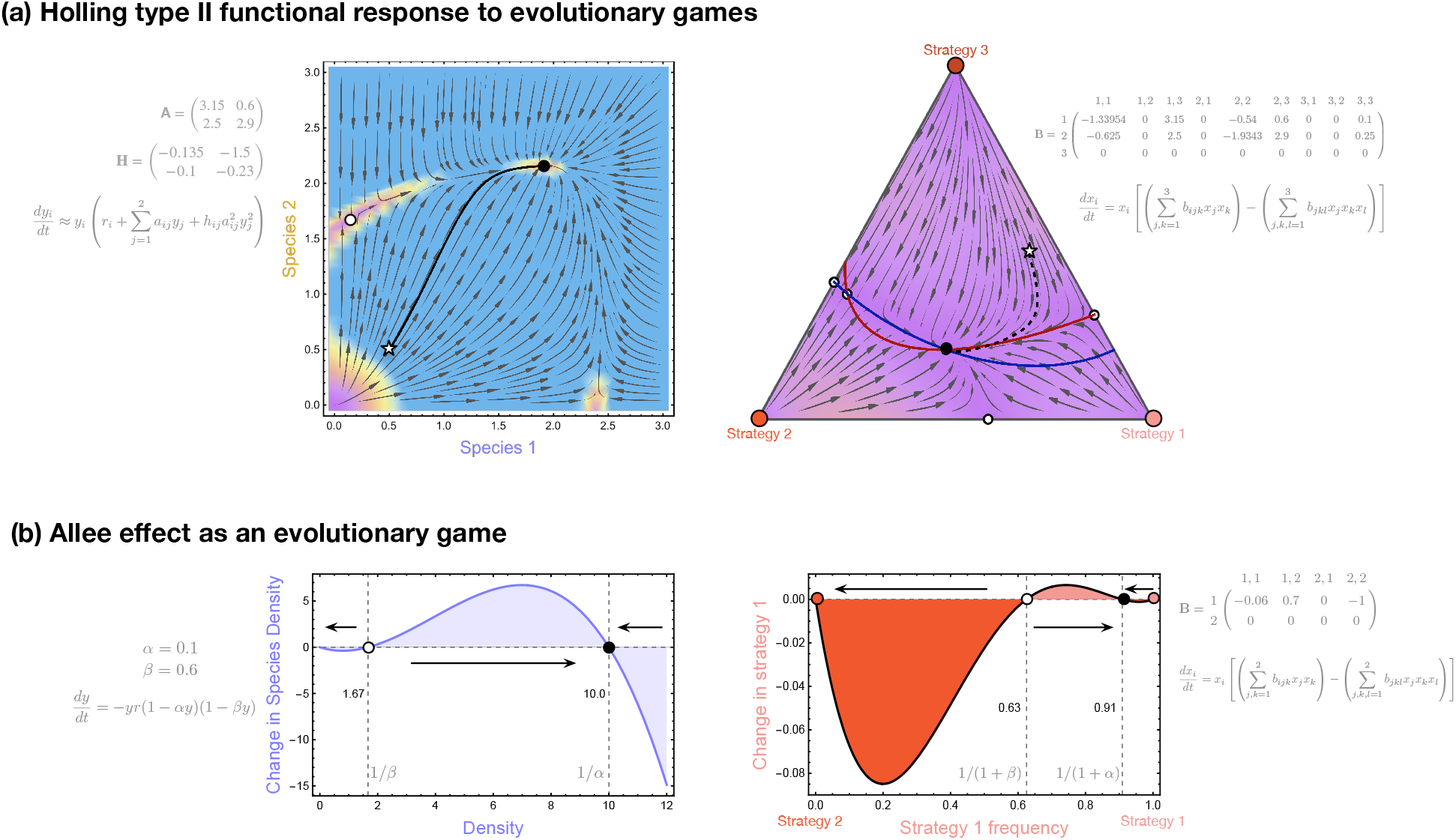
Equivalence between classic non-linear ecological models and multiplayer evolutionary games. **(a)** The two species interact with each other using a functional form as shown in (S13) up to the second order with interactions and handling times given by **A** and **H**. An equivalent replicator dynamics for a three-player game with three strategies is derived, maintaining the dynamical pattern with two internal fixed points. The payoff matrix **B** contains the three rows each for strategy *i* = 1, 2, 3 where the columns are the corresponding payoffs when the other two players have strategy *j, k*, going from 1 to 3 **(b)** The strong Allee effect, as shown on the left using (S14) with the internal equilibria at 1 /*β* and 1/ *α*. The population shows a negative growth rate for densities below 1 /*β*. It is only positive if the population density is between 1 /*β* and 1 /*α*. Above 1 /*α* the population suffers from overcrowding and declines. An equivalent three-player two-strategy replicator system shows the internal equilibria at 1 /(1 + *β*) and 1/(1 + *α*). The game demonstrates dynamics that can be described by a stag-hunt game with thresholds (*31*). Codes for generating the figures are available on Zenodo.

#### Allee effect

The Allee effect is crucial for understanding population dynamics, particularly in conservation and species management. The Allee effect is an ecological concept describing a phenomenon in which the overall population growth rate is positively correlated with population size or density (*46*). In other words, individuals in smaller populations may have lower survival or reproductive success than those in larger populations. This is typically modelled by a cubic differential equation, *y* = −*y* (1 – *αy*) (1 – *βy*). This effect can lead to critical thresholds, below which populations may decline toward extinction. The strong Allee effect is observed when a population must exceed a certain size or density before achieving a positive growth rate. Below this threshold, the population may continue to decline until it becomes extinct. Under the weak Allee effect, the population growth rate increases with population size, but the population can still grow even at very low densities. However, the population’s growth rate is lower at smaller sizes. Figure 3 bottom illustrates the strong Allee effect. The population exhibits negative growth at low densities. On the one hand, as per Allee’s arguments, organisms need each other to reproduce at low densities, even when resources are abundant, hence the existence of a lower threshold (*46*). On the other hand, high densities put pressure on resources and lead to population collapse. The corresponding three-player, two-strategy evolutionary game shows the two internal equilibria at 1 /(1 + *β*) and 1 / (1 + *α*). The lower equilibrium 1 / (1 + *β*) is unstable and serves as the Allee threshold: below it, the strategy frequency collapses; above it, it grows. The upper equilibrium1 /(1 + *α*)is stable and corresponds to the carrying-capacity bound. The resulting dynamics is a stag hunt with a threshold (*31*): a minimum number of hunters is required to secure the hunt (the lower threshold), and once this threshold is met, additional hunters provide no further benefit (saturation at the upper equilibrium).

## Discussion

While the equivalence between the replicator and the Lotka-Volterra systems is well-established for two-player, linear interactions between species interaction patterns (*6, 11*), the connection between these frameworks is often underappreciated, particularly when considering the dimensionality reduction from the replicator equations to the Lotka-Volterra systems (*14*). Here, we demonstrate the transfer of results from multiplayer evolutionary games to higher-order interactions in ecology via a transformation similar to that used for two-player games. This connection allows us to reinterpret results from evolutionary games within an ecological context, enabling us to leverage insights from evolutionary games in ecology – and vice versa.

The analysis is frequently numerical, and while excluding higher-order interactions can lead to the omission of crucial elements of observed biodiversity, the relationship between pairwise and higher-order interactions does not always guarantee stable or persistent coexistence. Specific choices between pairwise and higher-order interactions, such as “constrained higher-order interactions”, allow for the coexistence of multiple species, and although these choices are analytically challenging, they are accessible (*17*). Another approach to understanding complex ecological systems is to transform them into a well-studied replicator system (*11*). By formalizing the non-linear, higher-order connection, we have emphasized that while the transformation is mathematically useful, it does not imply a conceptual equivalence of interpreting strategies as species (*12*). For example, while intransitive interactions can be captured very well with both the ecological and evolutionary game theoretical models, a rock-paper-scissors relationship in ecology differs from a formal rock-paper-scissors game theoretical model.

The relationship between the replicator and the Lotka-Volterra equations is well-known for pairwise interactions, but it is not implemented in practical applications. Studies of non-linear evolutionary games with diverse group sizes (*47*) and the use of analytical methods such as Bernstein polynomials (*34*) enable the study of ecologically complex systems through the lens of evolutionary game theory – but until now, their translation to classical ecological models remained elusive. In infinitely large populations, research on the number and accessibility of equilibria provides insights into the potential for coexistence in evolutionary games (*48*). Extensions to multiplayer games increase the potential for internal equilibria (*49, 50*). Focusing on finite populations also allows for estimating coexistence probabilities and the long-term abundances of different strategies (*51*) and could open a new tool-set when translated into ecological terms.

Conversely, studies on higher-order interactions in ecology have had a strong empirical foundation (*44*). Only recently, the challenges of experiments with higher order interactions were overcome: Some experimental systems can now capture and quantify these intricate dynamics (*16, 52, 53*). Higher-order interactions in ecology naturally incorporate complexities such as founder effects or the impact of pioneer species, which may mediate interactions differently among the same group of interactors. For instance, the impact of species *j* on species *i*, when mediated via species *k*, can differ from the impact of species *k* on species *i* when mediated via species *j* (*14*). While multiplayer evolutionary games can model these situations effectively, justifying the differences in payoffs (e.g., between *b*_*ijk*_ and *b*_*ik j*_) does not always warrant the added complexity. Evolutionary games could thus benefit from interpretations informed by ecologically relevant scenarios and perhaps explore even novel game dynamics.

The equivalence between the replicator and the Lotka-Volterra equations is more general than previously known, but with two qualifications. First, the construction applies to polynomial non-linearities, Non-polynomial functional forms must either be approximated by a truncated polynomial expansion or recast through an alternative scheme such as a quasipolynomial embedding (*54, 55*). Second, this generality introduces increased complexity in interpreting interactions and correspondence between species and strategies. Already, in simpler cases, the correspondence between species and strategies is often lost during transformation (*11, 12*). While we can select and apply a specific transformation in both directions, alternative transformations are also possible. For example, one could define multiple payoff matrices for the same ecological scenarios. Although mathematically sound, this approach complicates the interpretation of interaction structures. In non-linear models,these interaction structures become even more complex (e.g., multiple interaction structures and payoff tensors as shown in Fig. 1), further muddling interpretations.

Given the challenges regarding terminology and timescales, understanding how different methodologies connect to corresponding Lotka-Volterra systems – and what they mean – will be a highly rewarding research program linking evolutionary games to rich ecological models, contributing further to a unification of models in theoretical biology.

## Acknowledgments

Funding from the Max Planck Society is graciously acknowledged. CSG thanks Prateek Verma and Frank Bastian for stimulating discussions. Feedback from the members of the Department of

Theoretical Biology, MPI Plön and the T-Eco-Evo group is appreciated.

## Author contributions

Both authors conceived the idea, performed the calculations, analysed the results, and wrote the manuscript.

## Data and materials availability

Figure-generating codes and the transformations between Lotka-Volterra and the replicator dynamics are available on Zenodo (https://doi.org/10.5281/zenodo.22677548).

## Supplementary materials

Materials and Methods

References (*1–57*)

## Supplementary Materials

### Materials and Methods

#### Transformation: pairwise interactions

##### From the Lotka-Volterra equations to the replicator dynamics

We begin with the Lotka-Volterra equations as denoted in Eqs. (1) rewritten below,

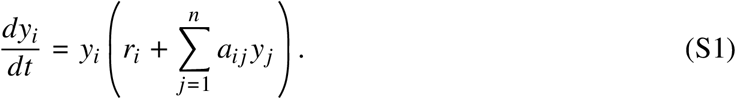

The set of equations are valid for *i* = 1, …, *n* species. To transform this set of *n* equations to a set of *n* + 1 replicator equations, we first introduce *y*_*n*+1_ = 1 and 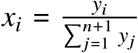 for *i* = 1, …, *n* + 1. The variables *x*_*i*_ with *i* = 1, … *n* + 1 lie on the simplex 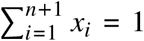. Using the above transformation, the Lotka-Volterra equations can be represented as

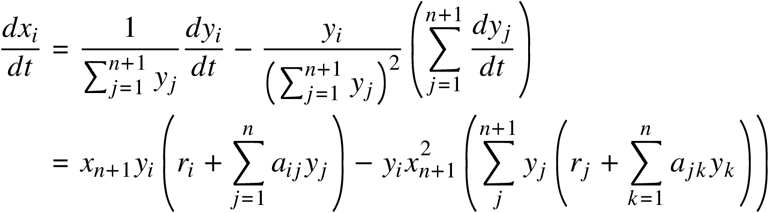

By subsuming the growth rates *r*_*i*_ of species *i* into an *n* +1 × *n* +1 matrix **B** at location *b*_*in*+1_ = *r*_*i*_ and setting *b*_*n*+1*i*_ = 0

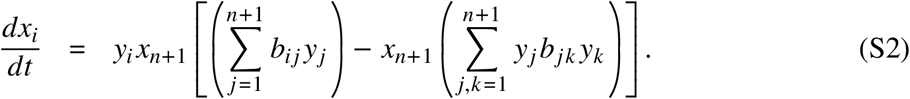

Using the inverse transformation of *y*_*i*_ = *x*_*i*_/*x*_*n*+1_ for *i* = 1, …, *n* we have

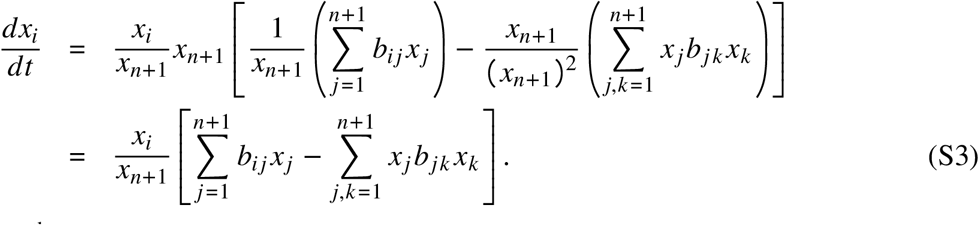

Note that the *n* + 1^th^ strategy then has no explicit interactions with the other *n* strategies as we have set *b*_*n*+1,*j*_ = 0 (*9, 56*). For all other entries, we have a direct correspondence between the species interaction matrix **A** and the payoff matrix **B** such that *b*_*i,j*_ = *a*_*i,j*_ for *i, j* from 1 to *n* (Fig. 1).

Dynamically rescaling time by a factor of *x*_*n*+1_ we have the replicator equations as in Eqs. (2),

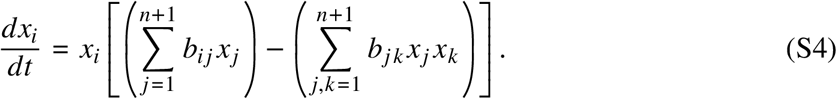

##### From the replicator dynamics to the Lotka-Volterra equations

To derive the reverse mapping from the replicator to the Lotka-Volterra dynamics we first invoke the inverse mapping from **x** ↦ **y**,

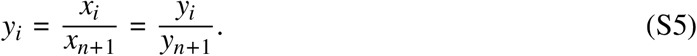

The derivative of *y*_*i*_ is

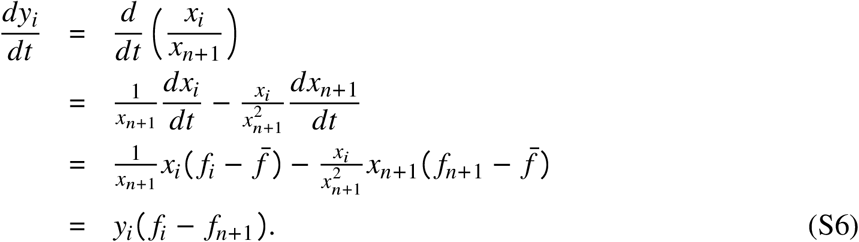

Note that we have not made any assumption about the particular form of *f*. For a two-player game, the payoffs are captured in a matrix **B** =[*b*_*ij*_], where *b*_*ij*_ denotes the payoff to strategy *i* when interaction with strategy *j*. The average fitness of strategy *i* is then denoted by 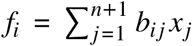.

Using this description, we have

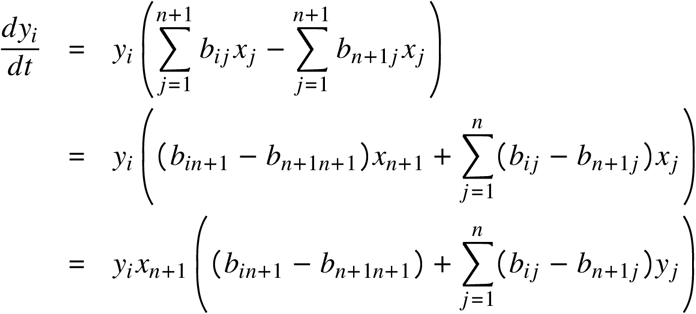

The differences in the payoffs can now be converted to the conventional notation used in the Lotka-Volterra dynamics, setting *r*_*i*_ = *b*_*in*+1_ − *b*_*n*+1*n*+1_ for the growth rates and *a*_*ij*_ = *b*_*ij*_ − *b*_*n*+1*j*_ for the interspecies interactions. Further, dynamically rescaling time by *x*_*n*+1_, we have the Lotka-Volterra equation as in Eqs. 1,

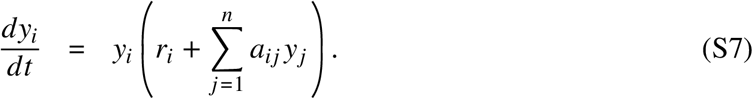

#### Transformation: Triplet interactions

##### From the Lotka-Volterra equations to the replicator dynamics

The interactions described in Eqs. (1) reflect the case where the focal species *i* interacts with another species *j* linearly. We can extend this interaction pattern to include higher-order interactions. A triplet Lotka-Volterra interaction is defined as an individual of a species interacting with two other individuals who may be from the same or different species, leading to a quadratic dependence. Combined with the pairwise interactions, the equation denoting the Lotka-Volterra dynamics is then given by

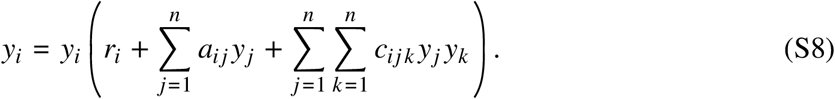

The interaction terms are of different order:

1. *r*_*i*_ is the constant growth rate of species *i*.
2. The matrix **A** with elements *a*_*ij*_ captures the pairwise effects of species *j* on species *i*, leading to a linear term.
3. The tensor **C** with elements *c*_*ijk*_ captures the effects of species *j* and *k* (in that order) on species *i*, leading to a quadratic term.

The payoff matrix for the corresponding evolutionary game is of the same dimension as the dimension of the highest order in the ecological scenario (in this case, three), but the number of strategies (*m*) increases by 1 (*m* = *n* + 1). Hence, the payoff tensor **B** is of size *m* × *m* × *m*. Similar to the inclusion of the *r*_*i*_ (single dimension) into the matrix (in the two-player case) **B** (two dimensions) in (S3), here, along with the growth rates, we also subsume the pairwise interactions into the tensor **B**. The effect of species *j* on species *i* appears in two places in the object **B**, *b*_*ijm*_ and *b*_*imj*_. Then we need to split *a*_*ij*_ into two terms at two locations *b*_*ijm*_ and *b*_*imj*_. How we split the value may depend on the exact question at hand to maintain any explanatory value of the model. For example, if the payoff is due to priority effects of action, then it may be possible that what *i* gets due to *j* is because of *j* acting on *i* or vice versa. Mathematically, the analysis is indifferent, and we can set one of the redundant values in the tensor **B** to 0 and the value of *a*_*ij*_ in one of the two slots. In this case, *m* acts as a dummy variable similar to the use of 0 in *a*_*i*0_ in Eq.(1.1) of the paper (*9*). By following the same logic as in Eqs. (S3), we can show that the equivalent replicator system for *m* = *n* + 1 strategies is then

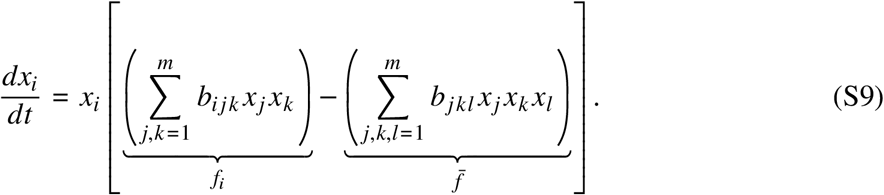

Similar to the case of pairwise interactions, the entries corresponding to the new strategy, *b*_*mjk*_ = 0 and the growth rates of the two species *r*_*i*_ appear at positions *b*_*imm*_.

##### From the replicator dynamics to the Lotka-Volterra equations

We now extend the logic from Appendix to a three-player game. A three-player game with *m* = *n*+1 strategies is denoted by **B** = *b*_*ijk*_. Each player can play one of the *m* strategies. For a three player game the average payoff for strategy *i* is then 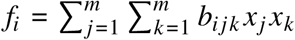.

Using (S6),

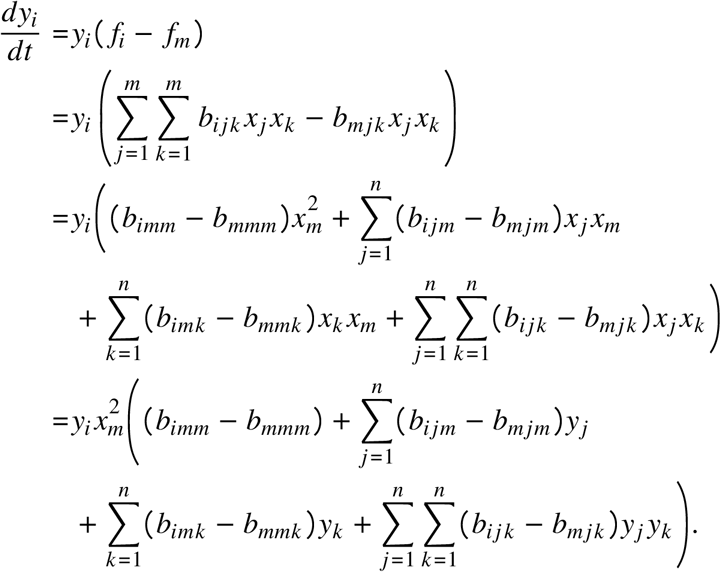

The *m*^*th*^ entries of **B** are being subtracted from the other entries of **B**. Recasting the payoff differences into terms usually used in the Lotka-Volterra dynamics, we have *r*_*i*_ = *b*_*imm*_ − *b*_*mmm*_. This is the intrinsic growth rate of species *i*. Next, since we now have a species *i* individual with two other individuals, we have the possibility that the ordering of the interactions matters. This combinatorial possibility is captured in the next two terms *c*_*ijn*_ = *b*_*ijm*_ − *b*_*mjm*_, *c*_*ink*_ = *b*_*imk*_ − *b*_*mmk*_. The final term including a double sum captures the general non-linear interaction where the densities of both the other interacting species matter simultaneously, *c*_*ijk*_ = *b*_*ijk*_ − *b*_*mjk*_. Dynamically rescaling time by 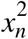, we get the Lotka-Volterra equivalent of a multiplayer game as,

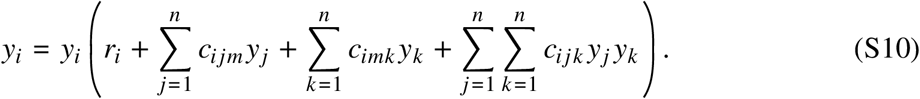

With this formulation, there is a single term of the order *O* (*y*^0^), which are the growth rates, two terms of order *O* (*y*), and one term of the order *O* (*y*^2^).

Note that here, in terms of degree 1, the index *m* appearing in the interaction parameters is a dummy variable. Again, this is similar to the use of 0 in *a*_*i*0_ in Eq.(1.1) of the paper (*9*). Allowing for a difference in *c*_*ijm*_ and *c*_*imj*_ allows for complete generality where priority effects or the impact of turn-based actions is important. Such interaction structures where the order matters can lead to rich dynamics and interesting results on the stability of the internal fixed point (*14, 33*). Simplification in the payoffs is possible, for example, if the player order does not matter. Hence, e.g. *c*_121_ = *c*_112_, the payoff to strategy 1 is the same whether the second player uses strategy 2 and the third uses 1 or vice versa. Then the two sums of degree 1 collapse into one, and we can drop the dummy variable leading to a simpler matrix capturing only pairwise interactions, **A** with entries *a*_*ij*_, such that

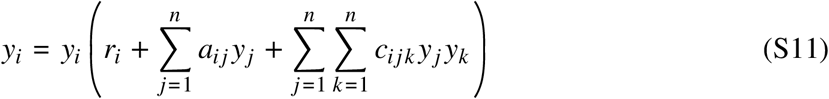

#### Transformation: general higher-order interactions

A general structure of a Lotka-Volterra dynamics that includes higher-order interactions is given by,

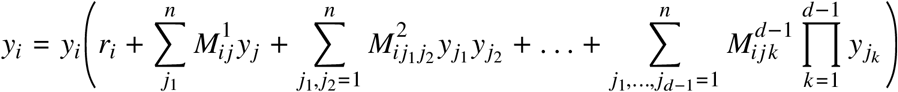

Here, the *r*_*i*_ capture the growth rates of the *i*th species. Each of the matrices *M*^*k*^ is of size *k* +1. Thus, *M*^1^ represents all the pairwise interactions while the tensors *M*^2^ collates all triplet interactions. We can extend this notation to include *d*^th^ order interactions captured in the tensor *M*^*d*−1^.

Generalising the algorithm from Appendix, an LV system with interactions up to order *d* – 1 maps to a *d*-player replicator system with *m* = *n* + 1 strategies, whose payoffs are stored in a *d*-dimensional tensor **B** of size *m*^*d*^. Write 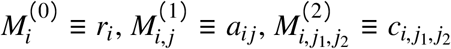, and in general let 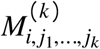denote the LV interaction coefficient of order *k*. Adopting the convention used throughout this paper – in which the dummy index *m* is packed into the leftmost positions and the species indices into the rightmost – the tensor entries are

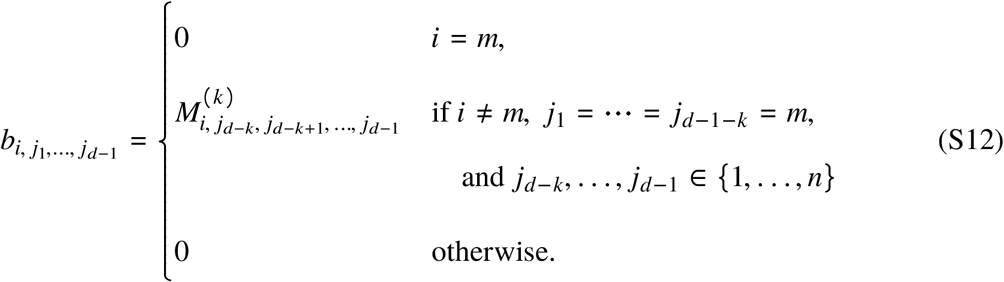

The species indices *j*_*d*−*k*_, …, *j*_*d*−1_ appearing as arguments of *M*^(*k*)^ on the right are the same variables appearing in the rightmost *k* slots of **B** on the left; the rule is that whatever species labels occupy those slots of **B** are passed, in the same order, as the species-label arguments of *M*^(*k*)^. The boundary cases of (S12) reproduce the constructions already given: *k* = 0 gives *b*_*i,m,m*,…,*m*_ = *r*_*i*_; *k* = 1 gives *b*_*i,m*,…,*m,j*_ = *a*_*ij*_ (matching *b*_*i,n*+1,*k*_ = *a*_*i,k*_ in the triplet case); *k* = *d* − 1 gives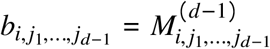. As discussed in the main text, this right-packed convention is one canonical representative within a many-to-one family: any redistribution of an order-*k* coefficient across its symmetric tensor placements — with placement weights summing to 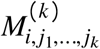 — yields the same LV system but, in general, a different multiplayer game. With **B** in hand, downstream analysis – deterministic or stochastic – can proceed with the standard machinery of multiplayer evolutionary games (*33, 51, 57*). Conversely, given a *d*-player game with *m* strategies, the construction inverted recovers an (*m* – 1)-species Lotka-Volterra system with interactions up to order *d* − 1, with time rescaled by a factor of 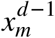.

Conversely, given a *d*-player game with *m* strategies, the same construction inverted recovers an (*m* − 1) - species Lotka-Volterra system with interactions up to order *d* − 1, with time rescaled by a factor of 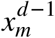.

While the transformation between Lotka-Volterra and the replicator equations (and back) works beautifully, we reiterate the often overlooked caveats in the discussion.

#### Deriving higher-order polynomials from other non-linearities

Here, we exemplify that we can obtain the higher-order terms needed for our transformation from other models.

##### Holling type II functional responses

First, we assume a Holling type II functional response. The Lotka-Volterra equations 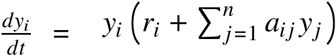change to

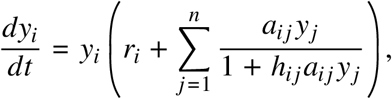

where *h*_*ij*_ can be interpreted as the time of species *i* interacting with species *j*, e.g. the handling time of a predator consuming a prey. If we assume that all interaction times are short, *h*_*ij*_ ≪ 1 for all *i* and all *j*, we obtain

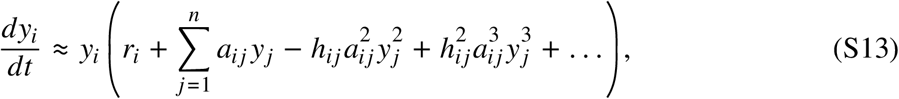

which has exactly the form that our transformation requires.

##### Holling type III functional responses

For Holling type III function responses, the dynamical equations change to

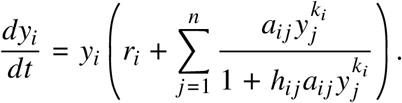

Small interaction times *h*_*ij*_ ≪ 1 lead to

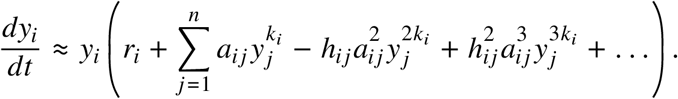

For integer *k*_*i*_ this has the desired polynomial form and the transformation given in Appendix applies directly. For non-integer *k*_*i*_ the equivalence does not extend as stated, since a *d*-player game requires an integer number of players. We anticipate that in practice, one must either approximate *k*_*i*_ by a rational number and rescale variables, or recast the system using a quasipolynomial embedding that introduces auxiliary species but reduces the dynamics to a standard polynomial form (*54*).

##### Allee effect

A version of the Allee effect is captured by the following population dynamical equation,

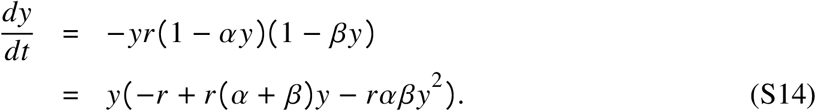

With *β* < *α*, we have a critical population density threshold 1 /*β* below which the population cannot survive. For population densities exceeding 1/ *β* but less than the carrying capacity threshold 1 / *α* the population grows. Exceeding the size 1 /*α*, the population then collapses. In the notation of Eq. (S8), we have *n* = 1 and thus only a single index, *i* = 1. Thus, *r*_1_ = −*r, a*_11_ = *r (α* + *β*), and *c*_111_ = −*rαβ*. Transforming them into our payoff tensor *b*_*ijk*_ with *i, j, k* = 1, 2, we have *b*_111_ = *c*_111_ = −*rαβ, b*_112_ = *r (α* + *β), b*_121_ = 0, *b*_122_ = −*r*, and *b*_211_ = *b*_211_ = *b*_221_ = *b*_222_ = 0. With this, we can utilize Eq. (S9) to convert to a three-player evolutionary game with two strategies,

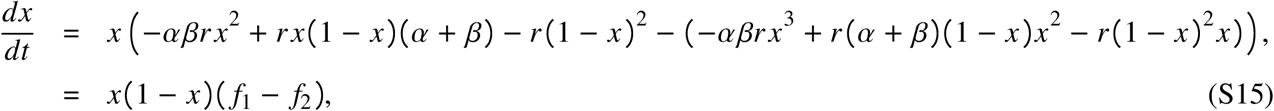

where *f*_1_ = −*r* + *r (*2 + *α* + *β) x* – *r(* 1 + *α* + *β* + *αβ) x*^2^ and *f*_2_ = 0 since the payoff entries for the second strategy are all set to 0. The two internal equilibria are given by 1/(1 + *α*) and 1/(1 + *β*).

